# Mouse parasubthalamic *Crh* neurons drive alcohol drinking escalation and behavioral disinhibition

**DOI:** 10.1101/2024.07.06.602357

**Authors:** Max Kreifeldt, Agbonlahor Okhuarobo, Jeffery L Dunning, Catherine Lopez, Giovana Macedo, Harpreet Sidhu, Candice Contet

## Abstract

Corticotropin-releasing factor (CRF, encoded by *Crh*) signaling is thought to play a critical role in the development of excessive alcohol drinking and the emotional and physical pain associated with alcohol withdrawal. Here, we investigated the parasubthalamic nucleus (PSTN) as a potential source of CRF relevant to the control of alcohol consumption, affect, and nociception in mice. We identified PSTN *Crh* neurons as a neuronal subpopulation that exerts a potent and unique influence on behavior by promoting not only alcohol but also saccharin drinking, while PSTN neurons are otherwise known to suppress consummatory behaviors. Furthermore, PSTN *Crh* neurons are causally implicated in the escalation of alcohol and saccharin intake produced by chronic intermittent ethanol (CIE) vapor inhalation, a mouse model of alcohol use disorder. In contrast to our predictions, the ability of PSTN *Crh* neurons to increase alcohol drinking is not mediated by CRF_1_ signaling. Moreover, the pattern of behavioral disinhibition and reduced nociception driven by their activation does not support a role of negative reinforcement as a motivational basis for the concomitant increase in alcohol drinking. Finally, silencing *Crh* expression in the PSTN slowed down the escalation of alcohol intake in mice exposed to CIE and accelerated their recovery from withdrawal-induced mechanical hyperalgesia. Altogether, our results suggest that PSTN *Crh* neurons may represent an important node in the brain circuitry linking alcohol use disorder with sweet liking and novelty seeking.

## Introduction

The neurocircuitry changes mediating the development and maintenance of alcohol use disorders are complex and dynamic (1). During withdrawal, the recruitment of stress-related signaling is thought to play a key role in the long-lasting dysregulation of hedonic homeostasis and the production of a negative emotional state fueling the escalation of alcohol use via negative reinforcement (2). Notably, release of the neuropeptide corticotropin-releasing factor (CRF, encoded by *Crh*) and subsequent activation of CRF_1_ receptors in the central nucleus of the amygdala (CeA) and bed nucleus of the stria terminalis (BNST) have been implicated in alcohol intake escalation and other behavioral indices of hyperkatifeia (e.g., negative affect, increased pain sensitivity) displayed by mice and rats withdrawn from chronic alcohol exposure (3–8). Identifying the relevant source of CRF, i.e., the cell body location of neurons releasing CRF during alcohol withdrawal, represents an important step toward the elucidation of the molecular and cellular mechanisms driving excessive alcohol drinking.

In rodent models of AUD combining alcohol self-administration with chronic intermittent exposure (CIE) to alcohol vapor inhalation (9, 10), CeA *Crh* neurons drive alcohol intake escalation in rats (11), but not in mice (12), even though mouse CeA *Crh* neurons are activated by and promote alcohol binge drinking (13–15). In the present study, we investigated the role of the parasubthalamic nucleus (PSTN) as a potential source of CRF that may be relevant to the behavioral adaptations induced by CIE in mice.

The PSTN is a small nucleus of the posterior lateral hypothalamus that sends dense projections to the CeA and BNST, as well as to other areas known to modulate alcohol drinking and the encoding of aversive experiences (16, 17). PSTN activity (as indexed by cFos expression) correlates more strongly with activity in amygdala nuclei in alcohol-drinking mice withdrawn from CIE than in air-exposed or alcohol-naïve counterparts (18). Furthermore, the PSTN contains a cluster of *Crh* cells and may thus serve as a source of CRF release in relevant brain regions during alcohol withdrawal. Accordingly, we used chemogenetics to test the hypothesis that PSTN *Crh* neurons can increase alcohol drinking, negative affect, and/or pain sensitivity in mice. We also measured saccharin drinking to evaluate whether the effects on alcohol intake generalize to another reinforcer. We then used local gene knockdown to test the hypothesis that *Crh* expression in the PSTN contributes to alcohol intake escalation and other behavioral phenotypes associated with CIE withdrawal.

## Methods

### Animals

C57BL/6J males were purchased from Scripps Research rodent breeding colony at 8 weeks of age. *Crh*-IRES-Cre (*Crh*-Cre) and *Tac1*-IRES2-Cre-D (*Tac1*-Cre) male breeders were obtained from The Jackson Laboratory (B6(Cg)Crh^tm1(cre)Zjh^/J, stock # 012704 (19); B6;129S-Tac1^tm1.1(cre)Hze/J^, stock #021877 (20)). Backcross breeders (C57BL/6J mice from Scripps Research rodent breeding colony) were introduced every 1-2 years to prevent genetic drift. All *Crh*-Cre and *Tac1*-Cre mice used for experimentation were heterozygous.

Mice were maintained on a 12 h/12 h light/dark cycle. Food (Teklad LM-485, Envigo) and reverse osmosis purified water were available *ad libitum* except for a 4-h period of water deprivation prior to water intake measurement and daily water restriction in the CNO-induced taste conditioning experiment. Sani-Chips (Envigo) were used for bedding substrate. All experiments included mice from both sexes, except for the CIE-2BC experiments, which used males only based on our experience of more robust alcohol intake escalation in this sex (21). Mice were at least 10 weeks old at the time of surgery and were single-housed throughout the duration of the experiments, starting at least 3 days prior to behavioral testing. All tests were conducted during the dark phase under red light unless otherwise specified.

All procedures adhered to the National Institutes of Health Guide for the Care and Use of Laboratory Animals and were approved by the Institutional Animal Care and Use Committee of The Scripps Research Institute.

### Viral vectors

Details of the viral vectors used in each experimental cohort are provided in Table S1. Adeno-associated viral (AAV) vectors encoding designer receptors for chemogenetic excitation (hM3Dq) or inhibition (hM4Di) (22, 23) fused to the red fluorescent protein mCherry, under the control of the human synapsin promoter and in a Cre-dependent manner, were obtained from the Vector Core at the University of North Carolina (UNC) at Chapel Hill or from Addgene (plasmids #44361 and #44362, respectively).

AAV vectors encoding a short hairpin RNA (shRNA) targeting the *Crh* mRNA (shCrh, 5’-GCATGGGTGAAGAATACTTCC-3’, selected using BLOCK-iT RNAi designer, loop sequence 5’-TTCAAGAGA-3’) or a control sequence (shControl, 5’-GTACGGTGAGCTGCGTTATCA-3’) under the control of a U6 promoter, along with a GFP reporter driven by a CBA promoter, were produced by Virovek. The shCrh and shControl constructs were packaged in an AAV8.2 capsid, in which the Phospholipase A2 domain encoded by the VP1 Cap gene is replaced with the corresponding domain from AAV2 to optimize endosomal escape (24, 25), by Virovek.

### Drugs

Clozapine-N-oxide (CNO) freebase was obtained from Enzo Life Sciences Inc. (BML-NS105-0025) or Hello Bio Inc. (HB1807), dissolved in dimethyl sulfoxide (DMSO), and diluted in 0.9% saline for intraperitoneal (i.p.) injection (10 mL/kg body weight) at a dose of 1 mg/kg (unless specified otherwise), 30 min prior to behavioral testing. The vehicle solution contained 0.5% DMSO.

CP376395 hydrochloride (Tocris Bioscience, 3212), riluzole (Tocris Bioscience, 0768), naloxone hydrochloride (MP Biomedicals, 0219024525), and acamprosate calcium (Tocris Bioscience, 3618) were dissolved in saline. MTEP hydrochloride (R&D Biosystems, 2921) was dissolved in Tween-80 and diluted in water (final Tween-80 concentration: 10%). JNJ16259685 (Tocris Bioscience, 2333) was dissolved in a vehicle made of 10% Captisol (w:v) in water. Aprepitant (Sigma, 1041904) was dissolved in DMSO and diluted in saline (final DMSO concentration: 1%). SR142948 (Tocris Bioscience, 2309) was dissolved in Tween-80 and diluted in saline (final Tween-80 concentration: 0.05%). Almorexant hydrochloride (Selleck Chemicals, S2160) was dissolved in a vehicle made of 20% Captisol (w:v) in water. SB222200 (Tocris Bioscience, 1393) was dissolved in a saline/0.3% Tween-80 vehicle. All these ligands were administered i.p., immediately prior to CNO/vehicle administration. Doses were selected based on publications reporting significant behavioral effects in mice (26–37).

Ethanol was obtained from PHARMCO-AAPER (200 proof for drinking and i.p. injection, 111000200; 95% for vaporization, 111000190). Pyrazole (Sigma-Aldrich, P56607) was dissolved in saline and administered i.p.

Chloral hydrate (Sigma-Aldrich, C8383) was dissolved in water at a concentration of 35% (w:v), and injected i.p.

### Experimental cohorts

Cohort details, including sample size by sex and corresponding figure panels, are provided in Table S1.

Three independent cohorts of *Crh*-Cre mice (Cohorts 1-3) were used to test the effect of chemogenetic activation of PSTN *Crh* neurons on fluid consumption (alcohol, saccharin, water) and affect (digging, tail suspension, elevated plus maze [EPM]). Cohorts 1-3 all included mice expressing hM3Dq; Cohort 2 also included a control group expressing mCherry only to control for potential off-target effects of CNO; Cohort 3 was injected with a smaller volume of viral vector to minimize viral transduction in adjacent brain regions. Cohort 1 was also used to test the ability of ligands to block the effect of CNO on alcohol drinking. Cohort 2 was also tested for nociception (tail pressure). The testing order was as follows: Cohort 1 – alcohol, saccharin, water, EPM, ligand testing; Cohort 2 – alcohol, saccharin, digging, tail suspension, EPM, tail pressure; Cohort 3 – water, saccharin, digging, alcohol, tail suspension. These tests were conducted on different weeks. The selection of tests and order of testing were designed to rule out potential carry-over effects and to perform the most stressful tests at the end. The effect of CNO was tested according to a within-subject design, except for EPM, which was conducted only once in each mouse. Except for CNO dose-responses, mice did not receive CNO more than once in any given week.

For ligand testing, mice were split into subgroups of equivalent alcohol intake on the two days preceding testing (Mon-Tue) and assigned to a ligand dose (each subgroup contained the same number of males and females); this dose was administered along with vehicle or CNO (within-subject design, counterbalanced order) on two consecutive days (Wed-Thu). The ligands were tested in the following order, with at least one week between ligands: CP376395, riluzole (30 mg/kg was initially included in the dose-response, but not administered beyond the first testing day due to major sedative effects precluding drinking behavior), aprepitant (10 mg/kg on one week, 25 mg/kg on the subsequent week), SR142948, naloxone, almorexant, acamprosate, MTEP, JNJ16259685.

The effect of chemogenetic inhibition of PSTN *Crh* neurons on alcohol and saccharin drinking was tested in a cohort of *Crh*-Cre mice exposed to CIE-2BC. Ethanol intake escalation was first established, then the hM4Di vector was injected, escalation was re-established, and the effect of CNO on alcohol drinking was tested. The mice were then switched to saccharin 2BC and the effect of CNO on saccharin drinking was tested during the second week.

A cohort of *Tac1*-Cre mice was used to test the effect of chemogenetic activation of PSTN *Tac1* neurons on alcohol and saccharin drinking.

*Crh* knockdown efficiency was first quantified by chromogenic *in situ* hybridization (CISH) in a group of alcohol-naïve C57BL/6J mice. The effect of *Crh* knockdown on alcohol drinking and CIE-induced phenotypes was then tested in a group of C57BL/6J mice exposed to CIE-2BC. These mice were first trained to drink alcohol and split into subgroups of equivalent baseline intake for assignment to the shControl or shCrh vector. 2BC sessions were resumed three weeks later and alternated with CIE every other week for five rounds to measure ethanol intake escalation. Mice were exposed to an additional week of CIE and tested in the EPM and digging assay 6 and 10 days later, respectively. They were exposed to a final week of CIE and tested in the tail pressure test 3 and 13 days later, and in the tail suspension test 6 days into withdrawal.

### Stereotaxic surgeries

Mice were anesthetized with isoflurane and placed in a stereotaxic frame (David Kopf Instruments, model 940). A small hole was drilled in the skull (David Kopf Instruments, 1474) and a 75-1000 nL volume of viral vector was injected bilaterally into the PSTN (AP −2.3 mm from bregma, ML ± 1.0-1.2 mm from the midline, DV −5.0-5.2 mm from the skull) using either a dual syringe pump (Harvard Apparatus) controlling the plungers of 10-μL Hamilton syringes connected to 33-gauge single injectors projecting 5 mm beyond a 26-gauge double guide cannula (Plastics One), or a microinjector pump (World Precision Instruments, UMP3T-2) with an attached 10-μL NanoFil syringe (World Precision Instruments) fitted with a 33-gauge NanoFil blunted tip (World Precision Instruments, NF33BL). The vector was infused at a rate of 0.1 μL/min for 10 min and the injectors were left in place for an additional 5-10 min to minimize backflow. The scalp was sutured using surgical thread. Mice were left undisturbed for at least one week post-surgery, and at least 4 weeks elapsed until the effect of CNO was tested.

### Alcohol drinking

Food pellets were placed in the bedding instead of the food hopper throughout the duration of the drinking experiments. Two-bottle choice (2BC) drinking sessions were conducted Mon-Fri, starting at the beginning of the dark phase and lasting 2 h. During these sessions, the home cage water bottle was replaced with two 50-mL conical tubes fitted with a rubber stopper and sipper tube assembly and filled with acidified water or ethanol 15% (v:v), respectively. The positions of the water and ethanol bottles were alternated every day and bottles were weighed at the end of each session. Bottles were also placed in an empty cage to generate spill control values that were subtracted from the weights lost in the alcohol and water bottles of experimental cages. Selectivity was calculated by dividing the weight of alcohol solution consumed by the total weight of fluids consumed during the session (alcohol + water) and multiplying by 100. Body weights were measured on a weekly basis to calculate ethanol intake (g/kg).

### Alcohol vapor inhalation

Alcohol intake escalation was induced by alternating weeks of voluntary alcohol drinking during limited-access 2BC sessions (described above) with weeks of forced chronic intermittent ethanol (CIE) exposure via vapor inhalation (38). During CIE weeks, CIE-2BC mice were exposed to 4 cycles (Mon-Fri) of 16 h ethanol vapor inhalation/8-h air inhalation followed by 72 h withdrawal (Fri-Mon). Ethanol was dripped into a heated flask using a metering pump (Walchem, EWN-B11PEUR), and an air pump (Hakko, HK-40LP) conveyed vaporized ethanol into custom chambers (modified from Allentown Sealed Positive Pressure individually ventilated cages). Mice received an i.p. injection of ethanol (1.5 g/kg) and pyrazole (68 mg/kg) before each 16-h ethanol vapor inhalation session. Blood alcohol levels (BALs) were measured on a weekly basis using gas chromatography and flame ionization detection (Agilent 7820A). The drip rate was adjusted to yield target BALs of 150-250 mg/dL. Control mice (Air-2BC) breathed air only and received pyrazole.

### Saccharin drinking

Saccharin (Sigma-Aldrich, S1002) was dissolved in drinking water at a 0.02% (w:v) concentration. 2BC drinking sessions were conducted as described above for alcohol. Saccharin intake is expressed as mg saccharin per kg body weight.

### Water intake

The home cage water bottle was removed for the first 4 hours of the dark phase to stimulate higher levels of water consumption during the test. A single bottle of water (same drinking tube design as described above for 2BC) was provided during the 2-h session.

### Digging test

The mouse was placed in a new, clean cage with a bedding thickness of 5 cm and no lid, and allowed to freely dig for 5 min (Fig. 5B) or 3 min (Fig. S6A). The latency to dig, number of digging bouts, and total digging duration were recorded. Testing was conducted under dim white light.

### Tail suspension test

The mice were suspended by their tails using adhesive tape wrapped around the tail approximately 2 cm from the tip and affixed to shelving. Prior to taping, the tail was inserted in a clear hollow cylinder (3.5-cm length, 1-cm diameter, 1 g) to prevent tail climbing behavior. The test lasted 6 min and the total duration of immobility was recorded.

### Elevated plus-maze

The apparatus consisted of two opposite open arms (30 cm length × 5 cm width), with a 0.3 cm lip, and two enclosed arms of the same size, with 15 cm high walls. The runways were made of gray (Fig. 5D and S7B-D) or black (Fig. S6C) acrylic and elevated 30 cm above the ground. The lips and walls were made of translucent acrylic. The end of the open arms (starting 5 cm away from the edge) was defined as the distal zone (the proximal zone represents the remainder). Testing began by placing an animal on the central platform of the maze facing an open arm. The test lasted 5 min and the maze was cleaned between subjects. The following measures were recorded using the ANY-maze (Stoelting Co.) behavioral tracking system: total distance traveled, time spent and number of entries in the closed arms, open arms proximal zone, and open arms distal zone.

### Tail pressure test

Mechanical nociceptive thresholds were assessed by applying pressure on the tail using a digital Randall-Selitto apparatus (Harvard Apparatus). The mice were first habituated to enter a restrainer pouch made of woven wire (stainless steel 304L 200 mesh, Shanghai YiKai) over three days. On testing days, the mouse was gently introduced into the restrainer and the distal portion of the tail was positioned under the conical tip of the apparatus. The foot switch was then depressed to apply uniformly increasing pressure onto the tail until the first nociceptive response (struggling or squeaking) occurred. The force (in g) eliciting the nociceptive response was recorded. A cutoff force of 600 g was enforced to prevent tissue damage. The measure was repeated on the medial and proximal parts of the tail of the same mouse, with at least 30 s between each measure. The average of the three measures (distal, medial, proximal) was used for statistical analysis.

### Splash test

A solution of 10% sucrose was sprayed on the dorsal coat of the mouse using a single squirt from a standard gardening spray bottle in mist position. The latency to groom, the number of grooming bouts, and duration of grooming were recorded during 5 min.

### Histology

At the end of all experiments, brains were analyzed to evaluate stereotaxic targeting accuracy by visualizing the mCherry and GFP reporters using native fluorescence or immunolabeling. Mistargeted mice were excluded from behavioral datasets accordingly (sample sizes reported in Table S1 include well-targeted mice only).

For native fluorescence and immunolabeling, the mice were anesthetized with chloral hydrate and perfused with cold PBS followed by 3.7% paraformaldehyde (PFA). Brains were dissected and immersion fixed in PFA for 2 hours at 4°C, cryoprotected in 30% sucrose in PBS at 4°C until brains sank, flash frozen in isopentane chilled on a dry ice ethanol slurry and stored at −80°C. Coronal 35-µm thick brain sections were sliced with a cryostat (Leica CM1950), collected in five series spanning the PSTN in PBS containing 0.01% sodium azide, and stored at 4°C.

For native fluorescence, sections were washed in PBS, plated on Superfrost plus glass slides (Fisher Scientific, 1255015), and air-dried. Coverslips were mounted using DAPI-containing Vectashield Hardset medium (Vector Laboratories, H1500). Images were captured using a Keyence BZ-X700 fluorescence microscope.

For immunolabeling, the sections were first blocked in PBS containing 0.3% Triton-X100, 1 mg/mL BSA, and 5% normal goat serum (NGS) for 1 h, then incubated with the primary antibody diluted in PBS containing 0.5% Tween-20 and 5% NGS (rabbit anti-mCherry antibody, Abcam, ab167453, RRID:AB_2571870, 1:5,000; chicken anti-mCherry antibody, Abcam, ab205402, RRID AB_2722769, 1:5,000; chicken anti-GFP antibody, Abcam, ab13970, RRID:AB_300798, 1:2000) overnight at 4°C. Following washes in PBS, sections were incubated with the secondary antibody diluted in PBS (goat anti-rabbit conjugated to Alexa Fluor 568, Life Technologies, A11004, RRID:AB_2534072, 1:500; goat anti-chicken conjugated to Alexa Fluor 488, Life Technologies, A11039, RRID:AB_142924, 1:500) for 2 h at room temperature, washed in PBS, and mounted and imaged as described above.

To quantify *Crh* knockdown, brains were subjected to CISH. Mice were quickly decapitated, and brains were snap-frozen in isopentane. Ten series of 20-μm coronal sections were sliced in a cryostat, directly mounted on Superfrost slides, and stored at −80°C. A pBlueScript plasmid containing the rat *Crh* cDNA (1.1 kb) was donated by Dr. Kelly Mayo (Northwestern University, Evanston, IL). Digoxigenin (DIG)-labeled riboprobes were synthesized using a kit (Roche 11277073910). Sections were post-fixed in PFA 4%, and then acetylated in 0.1 M triethanolamine pH 8.0, acetic acid 0.2%. Following washes in salt sodium citrate (SSC) 2x, sections were dehydrated and defatted in a graded ethanol/chloroform series. Pre-hybridization and hybridization were performed at 70°C in a buffer containing 50% formamide, SSC 2x, Ficoll 0.1%, polyvinylpyrrolidone 0.1%, bovine serum albumin 0.1%, sheared salmon sperm DNA (0.5 mg/mL) and yeast RNA (0.25 mg/mL). Probes were diluted in the hybridization buffer (800 ng/mL) and incubated overnight on slides. Post-hybridization washes were performed in 50% formamide, SSC 2x, and Tween-20 0.1%. Sections were then blocked for 1 h and incubated with an anti-DIG antibody (Roche 11093274910, RRID:AB_514497, 1:2000) diluted in MABT buffer (0.1 M maleic acid pH 7.5, 0.15 M NaCl, Tween-20 0.1%) containing 10% NGS overnight at 4°C. Following washes in MABT and incubation in detection buffer (0.1 M Tris-HCl pH 9.5, 0.1 M NaCl, 0.05 M MgCl2, Tween-20 0.1%), the reaction with NBT-BCIP was allowed to develop in the dark for 24 h at room temperature. Slides were rinsed, air dried and mounted in DPX (Sigma-Aldrich). Sections containing the PSTN were imaged using a Zeiss Axiophot microscope equipped with a QImaging Retiga 2000R color digital camera and QCapture software. Images were converted to grayscale and optical density of the CISH signal in the PSTN was analyzed using NIH Image J software.

The co-localization of *Crh* with other genes was assessed in naïve C57BL/6J PSTN sections using the RNAscope Fluorescent Multiplex manual assay (ACD, 320851). Mice were perfused with cold PBS, quickly decapitated, and brains were snap-frozen in isopentane. Ten series of 20-μm coronal sections were sliced in a cryostat, directly mounted on Superfrost slides, and stored at −80°C. The kit protocol was followed (ACD documents 320513 and 320293), except that Protease III was used in lieu of Protease IV, slides were covered with Rinzl plastic coverslips during incubation steps, and probes were hybridized for 3 h. The following probes were used: mouse Crh (316091-C2), mouse Gad2 (439371), mouse Slc17a6 (319171-C3), mouse Penk (318761), mouse Tac1 (410351), and mouse Nts (420441). Sections containing the PSTN were imaged using a Zeiss Axiophot microscope equipped with a QImaging Retiga 2000R color digital camera and QCapture software. PSTN cells containing signal for each probe were counted and the percentage of colocalization with *Crh* was calculated.

### Statistical analysis

Data analysis was performed in GraphPad Prism (v10.2.3). The effects of CNO on behavioral measures were analyzed by paired t-test or by repeated-measures two-way analysis of variance (ANOVA) with sex, vector, ligand dose, or vapor exposure as between-subjects factor.

Significant interactions were followed by Dunnett’s multiple comparisons to Vehicle for dose-responses, and Šídák’s multiple comparisons otherwise. EPM data from each zone were analyzed by two-way ANOVA, followed when relevant by Šídák’s multiple comparisons. *Crh* CISH data were analyzed by unpaired t-test. The effects of *Crh* knockdown and vapor exposure on weekly alcohol intake and other behavioral measures were analyzed by two-way ANOVAs; unprotected Fisher’s Least Significant Difference tests were also used to examine the effect of CIE among shControl and shCrh mice independently (signaled by red stars). BALs were analyzed by two-way repeated-measures ANOVA. All t-tests were two-tailed. For repeated-measures ANOVAs, the Geisser-Greenhouse correction was used. Mice were excluded from a given dataset if their ethanol (saccharin) intake in the vehicle condition was lower than 0.3 g/kg (0.3 mg/kg, respectively), or if their value met with the Grubbs’ outlier criterion (no more than 1 mouse excluded per experimental subgroup) (39). In graphs, individual values and group averages are plotted and the error bars represent the standard error of the mean.

## Results

### Chemogenetic stimulation of PSTN *Crh* neurons promotes the consumption of alcohol and saccharin but not water

To test whether PSTN *Crh* neurons influence the voluntary consumption of alcohol, *Crh*-Cre mice were injected in the PSTN with a Cre-dependent vector encoding the excitatory designer receptor hM3Dq and trained to consume alcohol in limited-access, free-choice sessions (2-h 2BC, Fig. 1A). Chemogenetic stimulation of PSTN *Crh* neurons produced a robust increase in ethanol intake (Fig. 1B). This effect was significant across the three doses of CNO tested (main effect of dose: F_2.3,39.2_=23.3, p<0.0001) and in both sexes (sex x dose interaction: F_3,51_=1.3, p=0.28), although females consumed more alcohol than males overall (main effect of sex: F_1,17_=9.60, p=0.0065). There was no significant effect of CNO or sex on water intake (Fig. S1A) or selectivity (Fig. S1B) during 2BC sessions (F’s < 1.0, p’s > 0.50).

**Figure 1.**
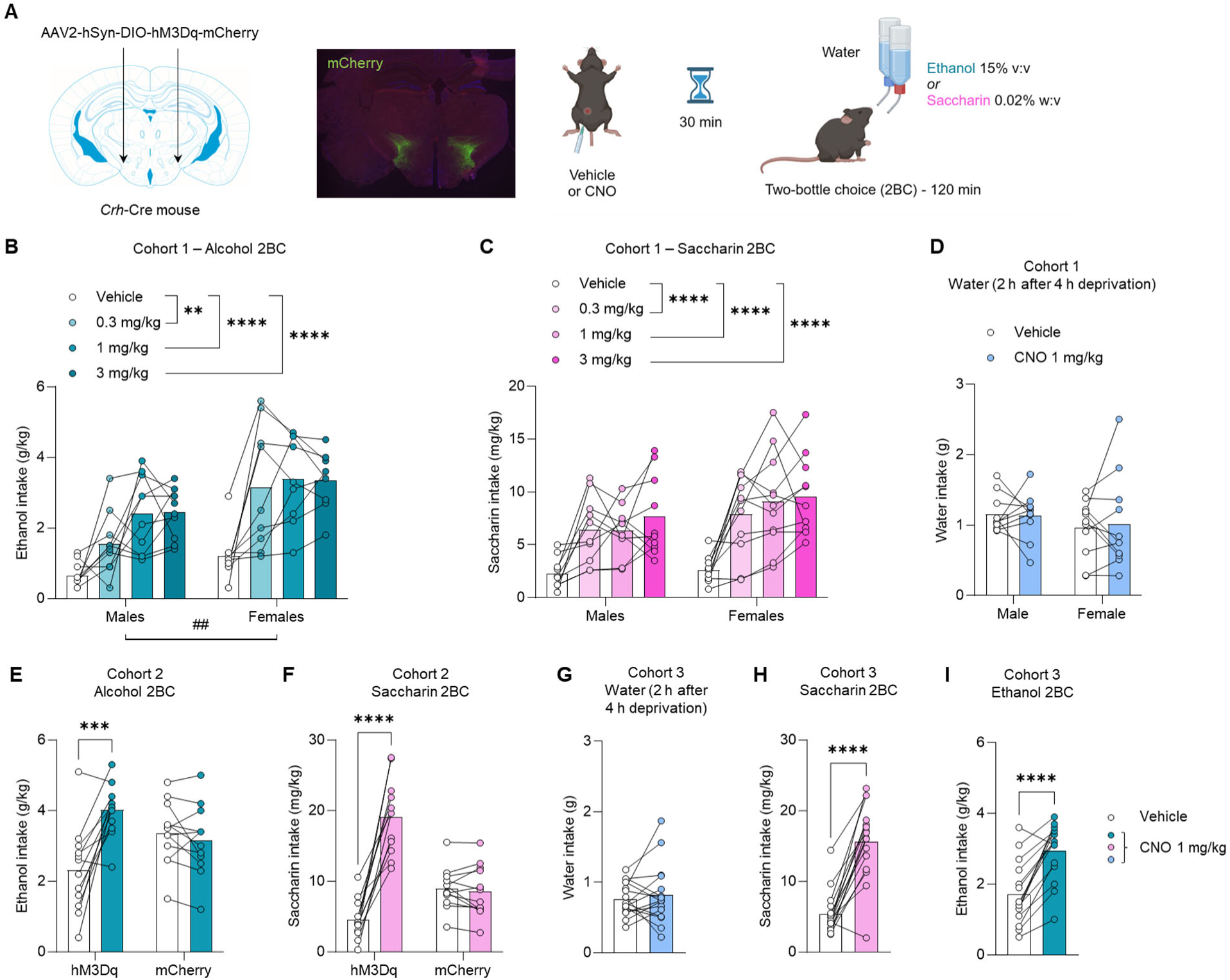
Chemogenetic stimulation of PSTN *Crh* neurons promotes the consumption of alcohol and saccharin but not water. **A.** *Crh*-Cre mice were injected with a Cre-dependent hM3Dq-encoding vector (or mCherry control) in the PSTN. mCherry immunolabeling shows targeted transduction in the PSTN. Voluntary alcohol (**B, E, I**) and saccharin (**C, F, H**) consumption was measured in 2-h two-bottle choice (2BC) sessions started 30 min after CNO injection. Water intake (**D, G**) was measured during 2 h after 4 h of water deprivation. Three independent cohorts were tested (Cohort 1, **B-D**; Cohort 2, **E-F**; Cohort 3, **G-I**). **B-C.** CNO vs. Vehicle: **, p<0.01; ****, p<0.0001, Dunnett’s *posthoc* tests. Main effect of sex: ^##^, p<0.01. **E-F.** CNO vs. Vehicle: ***, p<0.001; ****, p<0.0001, Šídák’s *posthoc* tests. As expected, CNO had no significant effect in mice expressing mCherry only. **G-I.** CNO vs. Vehicle: ****, p<0.0001, paired t-test.

The mice were then trained to consume saccharin in the same protocol (2-h 2BC). Chemogenetic stimulation of PSTN *Crh* neurons produced a robust increase in saccharin intake (Fig. 1C), across CNO doses (F_2.6,49.9_=26.1, p<0.0001) and sexes (main effect: F_1,19_=2.3, p=0.15; interaction: F_3,57_=0.88, p=0.46). There was a significant main effect of CNO on water intake (Fig. S1C; F_2.0,37.2_=3.8, p=0.031), reflecting an increase in water consumption in mice injected with 1 mg/kg (p=0.037) or 3 mg/kg (p=0.055) CNO. There was also a significant main effect of CNO on selectivity (Fig. S1D; F_1.9,36.9_=4.7, p=0.016), reflecting an increase in saccharin preference in mice injected with 0.3 mg/kg (p=0.028) or 1 mg/kg (p=0.056) CNO. From that point onward, all subsequent chemogenetic experiments used a CNO dose of 1 mg/kg.

We then probed whether the increased consumption of alcohol and saccharin might result from thirst, such that the chemogenetic stimulation of PSTN *Crh* neurons would also promote the consumption of water when offered as the sole available fluid. For this experiment, the water bottle was removed from the home cage for 4 h at the beginning of the dark phase and replaced with a water-containing drinking tube for the following 2 h. This mild water restriction was designed to produce water consumption levels similar to the volumes of alcohol or saccharin solution consumed during 2BC sessions. CNO did not significantly affect water intake (Fig. 1D; main effect of CNO: F_1,19_=0.025, p=0.88; main effect of sex: F_1,19_=0.85, p=0.37; interaction: F_1,19_=0.14, p=0.71).

To verify that the effects of CNO were driven by hM3Dq activation, rather than by a non-selective action of CNO or its metabolites on endogenous targets, we generated another cohort of *Crh*-Cre mice that received a Cre-dependent vector encoding hM3Dq or the mCherry reporter alone. In alcohol 2BC (Fig. 1E), there was a significant vector x CNO interaction (F_1,22_=11.1, p=0.0031), whereby CNO increased ethanol intake solely in hM3Dq mice (p=0.0002) and had no effect in mCherry controls (p=0.84). There were no significant effects of CNO or vector on water intake (Fig. S2A) or alcohol preference (Fig. S2B) (F’s<0.4, p’s>0.50). In saccharin 2BC (Fig. 1F), there was also a significant vector x CNO interaction (F_1,22_=66.8, p<0.0001), whereby CNO increased ethanol intake solely in hM3Dq mice (p<0.0001) and had no effect in mCherry controls (p=0.95). The CNO x vector interaction was trending for water intake during saccharin 2BC (Fig. S2C; F_1,20_=2.9, p=0.10), such that CNO increased water consumption in hM3Dq (p=0.0073) but not mCherry (p=0.63) mice. There was no significant effect of CNO or vector on saccharin preference (Fig. S2D) (F’s<2.0, p’s>0.18).

To rule out a potential role of testing order, whereby saccharin consumption might have been influenced by the prior consumption of alcohol, we generated a third cohort of *Crh*-Cre mice with Cre-dependent expression of hM3Dq in the PSTN. Consistent with the results obtained in the first cohort, CNO did not impact water intake when available as the sole fluid after a short period of deprivation (Fig. 1G; t_15_=0.68, p=0.51). The mice were then given access to saccharin 2BC. CNO again increased saccharin intake (Fig. 1H; t_15_=8.4, p<0.0001), but did not affect water intake (Fig. S3A; t_15_=1.5, p=0.14) or selectivity (Fig. S3B; t_15_=1.6, p=0.13). Alcohol 2BC was tested next. Likewise, CNO increased ethanol intake (Fig. 1I; t_14_=6.4, p<0.0001) without affecting water intake (Fig. S3C; t_15_=1.3, p=0.23) or selectivity (Fig. S3D; t_14_=0.70, p=0.50). Altogether, these data indicate that chemogenetic stimulation of PSTN *Crh* neurons stimulates the voluntary consumption of both alcohol and saccharin in male and female mice trained to drink these reinforcers, regardless of the order in which they are introduced. This effect does not result from a general increase in thirst as stimulating PSTN *Crh* neurons did not impact water intake in mildly deprived mice. The increase in water intake noted during saccharin 2BC may be related to taste dilution and was not as pronounced as the increase in saccharin intake. We also confirmed the selectivity of our chemogenetic approach as no effects of CNO were detected in mice that do not express hM3Dq.

### Chemogenetic inhibition of PSTN *Crh* neurons reduces alcohol and saccharin consumption in the CIE-2BC model

We next tested whether the endogenous activity of PSTN *Crh* neurons might contribute to ethanol intake escalation in the CIE-2BC mouse model. To do so, *Crh*-Cre mice were injected into the PSTN with a Cre-dependent vector encoding the inhibitory designer receptor hM4Di. They were trained to drink alcohol in 2-h 2BC sessions and exposed on alternated weeks to chronic intermittent ethanol vapor inhalation (CIE), a regimen that produces an increase in voluntary alcohol consumption in CIE mice compared to control counterparts inhaling air only (38) (Fig. 2A). Chemogenetic inhibition of PSTN *Crh* neurons reduced alcohol consumption in both groups (Fig. 2B; main effect of CIE: F_1,12_=8.4, p=0.014; main effect of CNO: F_1,12_=7.1, p=0.021). Although the CIE x CNO interaction did not reach significance (F_1,12_=1.8, p=0.20), there was a trend for CIE mice to respond more strongly to CNO than Air mice, and CNO-treated CIE-2BC mice reduced their intake to the level of vehicle-treated Air-2BC mice. There was no significant effect of CIE or CNO on water intake (Fig. S4A) or selectivity (Fig. S4B) (F’s < 0.4, p’s > 0.54).

**Figure 2.**
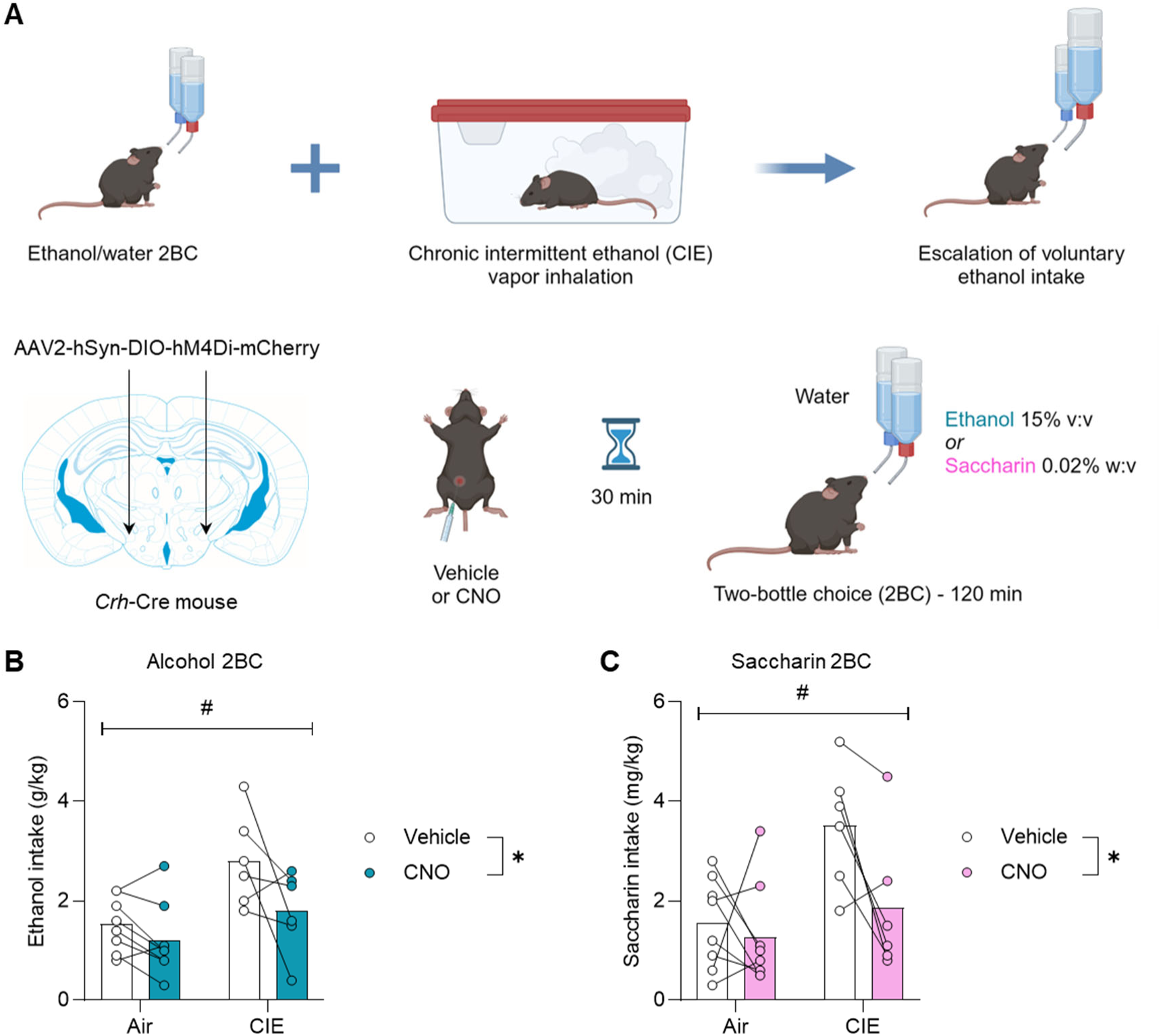
Chemogenetic inhibition of PSTN *Crh* neurons reduces alcohol and saccharin consumption in the CIE-2BC model. **A.** Mice were exposed to weeks of limited-access alcohol 2BC drinking alternated with weeks of chronic intermittent ethanol (CIE) vapor inhalation, a procedure that escalates voluntary alcohol consumption compared to air-exposed counterparts (Air). *Crh*-Cre mice were injected with a Cre-dependent hM4Di-encoding vector in the PSTN. Voluntary alcohol (**B**) and saccharin (**C**) consumption was measured following administration of CNO (within-subjects) 30 min prior to 2BC. Main effect of vapor: ^#^, p<0.05; main effect of CNO: *, p<0.05.

The mice were then given access to saccharin 2BC. The effects of CIE and CNO followed the same pattern as for alcohol 2BC. CIE mice consumed more saccharin than Air mice and the chemogenetic inhibition of PSTN *Crh* neurons reduced saccharin consumption in both groups (Fig. 2C; main effect of CIE: F_1,12_=7.6, p=0.018; main effect of CNO: F_1,12_=5.8, p=0.033).

Although the CIE x CNO interaction did not reach significance (F_1,12_=2.9, p=0.11), there was a trend for CIE mice to respond more strongly to CNO than Air mice, and CNO-treated CIE-2BC mice reduced their intake to the level of vehicle-treated Air-2BC mice. There was no significant effect of CIE or CNO on water intake (Fig. S4C) or selectivity (Fig. S4D) (F’s < 0.6, p’s > 0.44). Altogether, this data suggests that the endogenous activity of PSTN *Crh* neurons during withdrawal from CIE promotes alcohol and saccharin drinking.

### Alcohol consumption stimulated by PSTN *Crh* neuronal activation resists pharmacological inhibition

We then sought to address the signaling mechanism that mediates the increase in alcohol drinking driven by PSTN *Crh* neurons. Given the literature implicating CRF_1_ signaling in excessive alcohol drinking (3, 4, 6), we reasoned that blocking CRF_1_ receptors would prevent this effect. *Crh*-Cre mice expressing Cre-dependent hM3Dq in the PSTN and trained to drink alcohol were co-injected (i.p.) with CNO (or vehicle) and different doses of the CRF_1_ receptor antagonist CP376395 before 2BC (Fig. 3A). As expected, there was a significant main effect of CNO (F_1,21_=39.0, p<0.0001) reflecting the increase in alcohol drinking induced by chemogenetic stimulation of PSTN *Crh* neurons. However, the main effect of CP376395 (F_2,21_=0.5, p=0.61) was not significant. There was a trend for CNO x CP376395 interaction (F_2,21_=2.9, p=0.080) whereby the antagonist tended to reduce alcohol intake in vehicle-injected mice (p=0.069-0.076 vs. saline) but alcohol consumption following CNO administration was insensitive to CRF_1_ blockade (p=0.52-0.54). This finding indicates that the increased alcohol drinking produced by chemogenetic stimulation of PSTN *Crh* neurons does not require CRF_1_ signaling.

**Figure 3.**
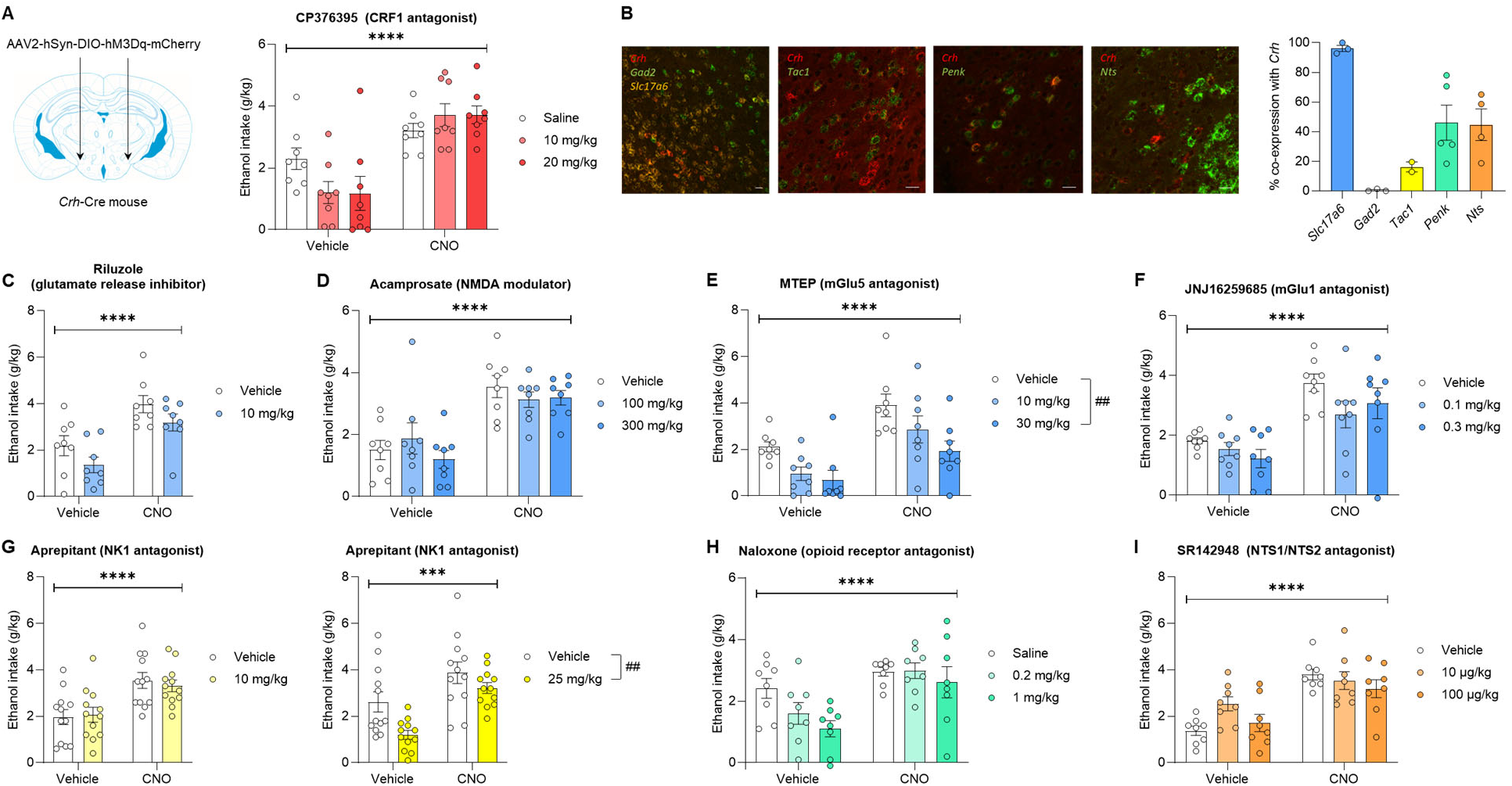
Alcohol consumption stimulated by PSTN *Crh* neuronal activation resists pharmacological inhibition. **A.** *Crh*-Cre mice were injected with a Cre-dependent hM3Dq-encoding vector in the PSTN and a CRF_1_ antagonist (CP376395) was administered at the same time as CNO (1 mg/kg) 30 min prior to alcohol 2BC. Ethanol intake was measured following combined administration of CP376395 (between-subjects) and CNO (within-subjects). Main effect of CNO: ****, p<0.0001. The CNO x CP376395 interaction trended toward significance (p=0.080). **B.** The cellular colocalization of *Crh* mRNA with markers of GABAergic (*Gad2*) or glutamatergic (*Slc17a6*) neurons, or with neuropeptide-encoding mRNAs known to be expressed in the PSTN (*Tac1*, *Penk*, *Nts*), was visualized by fluorescent *in situ* hybridization. Colocalization quantification shows that *Crh* is expressed by PSTN glutamatergic neurons and partially overlaps with the distribution of *Penk* and *Nts*, and to a small extent with *Tac1*. **C-I.** *Crh*-Cre mice were injected with a Cre-dependent hM3Dq-encoding vector in the PSTN. A ligand (between-subjects) was administered at the same time as CNO (1 mg/kg, within-subjects) 30 min prior to alcohol 2BC. Ligands were selected to target glutamatergic transmission (**C-F**), NK1 receptors putatively activated by *Tac1*-encoded peptides (**G**), opioid receptors putatively activated by *Penk*-encoded peptides (**H**), and NTS1/NTS2 receptors putatively activated by *Nts*-encoded neurotensin (**I**). The only ligands that significantly altered alcohol intake were MTEP (**E**) and aprepitant at 25 mg/kg (**G**). Main effect of CNO: ***, p<0.001; ****, p<0.0001. Dunnett’s *posthoc* test vs. Vehicle (**E**) or main effect of ligand (**G**): ^##^, p<0.01. The CNO x ligand interaction did not reach significance for any of the tested ligands.

We reasoned that other neurotransmitters or neuromodulators released by PSTN *Crh* neurons might contribute to the effect of their chemogenetic stimulation on alcohol drinking. We characterized the co-localization of *Crh* with markers of glutamatergic (*Scl17a6*) and GABAergic (*Gad2*) neurons, as well as neuropeptide-encoding transcripts expressed at high levels in the PSTN (Fig. 3B). Consistent with the neurochemical makeup of the PSTN (17), virtually all PSTN *Crh* neurons express *Slc17a6* and no *Gad2*. Consistent with a previous report (40), we observed a limited overlap (∼15%) between *Crh* and *Tac1*, the transcript encoding substance P. A larger fraction of PSTN *Crh* cells (∼40%) co-express *Penk* or *Nts*, the transcripts encoding enkephalins and neurotensin, respectively.

Following the same approach as in Fig. 3A, we then tested whether antagonizing glutamate, substance P, enkephalin, or neurotensin signaling would compromise the ability of CNO to increase voluntary alcohol drinking in *Crh*-Cre mice expressing Cre-dependent hM3Dq in the PSTN (Fig. 3C-I and Fig. S4A). We also tested the potential involvement of orexin signaling, as this neuropeptide is produced in the lateral hypothalamic area, adjacent to the PSTN, where *Crh* neurons also reside (Fig. S4B). For each target, the antagonist (or its vehicle) was co-injected with CNO (or vehicle) prior to alcohol 2BC. With all ligands tested, the main effect of CNO was highly significant (F’s>24.6, p’s<0.0001) and the CNO x antagonist interaction did not reach significance (F’s<2.5, p’s>0.10). There was a significant main effect of antagonist for only two of the ligands: MTEP, a metabotropic glutamate receptor 5 (mGlu5) antagonist (Fig. 3E; F_2,21_=6.4, p=0.0067; 10 mg/kg, p=0.060; 20 mg/kg, p=0.0038), and aprepitant, a neurokinin receptor 1 (NK1) antagonist, at 25 mg/kg (Fig. 3G; F_1,22_=10.3, p=0.0040). Both MTEP and aprepitant reduced alcohol intake without affecting water intake during 2BC (Fig. S5A-B; F_2,20_=0.80, p=0.46 and F_1,21_=0.13, p=0.72, respectively). With riluzole, a glutamate release inhibitor, we initially included a higher dose of 30 mg/kg, but it produced overt sedation, so we limited our analysis to the lower dose of 10 mg/kg, which tended to reduce alcohol intake (Fig. 3C; F_1,14_=3.3, p=0.09). Naloxone, an opioid receptor antagonist, produced a similar trend (Fig. 3H; F_2,21_=2.6, p=0.10). A trend for CNO x SR142948 interaction (F_2,21_=2.5, p=0.11) was driven by SR142948, an NTS1/NTS2 receptor antagonist, increasing baseline (p=0.027 vs. vehicle) but not CNO-induced (p=0.79) alcohol drinking at 10 μg/kg. This observation is consistent with NTS1 positive allosteric modulation reducing and NTS2 gene knockout increasing ethanol consumption (41, 42) and rules out neurotensin signaling as a mechanism promoting alcohol drinking following the activation of PSTN *Crh* neurons.

Overall, we found that inhibiting mGlu1 receptors (Fig. 3F), opioid receptors (Fig. 3H), NTS1/NTS2 receptors (Fig. 3I), NK3 receptors (Fig. S4A), or OX1/OX2 receptors (Fig. S4B) does not affect the excessive alcohol drinking driven by PSTN *Crh* neurons. Several of these drugs tended to reduce alcohol consumption after vehicle treatment but failed to exert the same effect after CNO treatment (JNJ16259685, naloxone, SB222200, almorexant). In contrast, blocking mGlu5 and NK1 receptors reduced alcohol drinking regardless of CNO vs. vehicle pretreatment. Even in the presence of mGlu5 and NK1 antagonists, ethanol intake following CNO injection remained higher than following vehicle injection, indicating that PSTN *Crh* neurons can at least partially overcome the suppressive effect of these ligands on alcohol drinking. In conclusion, alcohol consumption stimulated by the activation of PSTN *Crh* neurons resists pharmacological manipulations that reduce alcohol drinking under control conditions.

### Chemogenetic stimulation of PSTN *Tac1* neurons reduces the consumption of reinforcing fluids

Since we observed a significant reduction of alcohol intake by aprepitant and the main endogenous activator of NK1 receptors is substance P (encoded by *Tac1*), which is expressed in a large fraction of PSTN neurons, we tested whether PSTN *Tac1* neurons might promote alcohol drinking to the same extent as PSTN *Crh* neurons. To do so, *Tac1*-Cre mice were injected in the PSTN with a Cre-dependent hM3Dq vector and trained to drink alcohol in 2-h 2BC sessions (Fig. 4A). Chemogenetic stimulation of PSTN *Tac1* neurons strongly reduced ethanol intake (Fig. 4B; t_4_=7.1, p=0.0021). These mice were then trained to drink saccharin and CNO likewise strongly reduced saccharin intake (Fig. 4C; t_4_=6.6, p=0.0027).

**Figure 4.**
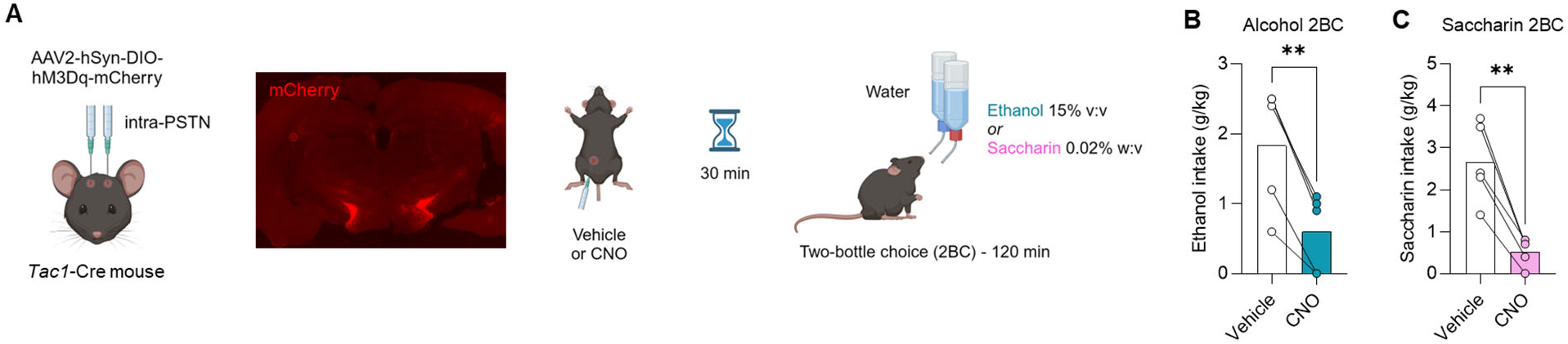
Chemogenetic stimulation of PSTN *Tac1* neurons reduces the consumption of reinforcing fluids. **A.** *Tac1*-Cre mice were injected with a Cre-dependent hM3Dq-encoding vector in the PSTN. mCherry fluorescence shows targeted transduction in the PSTN. Voluntary alcohol (**B**) and saccharin (**C**) consumption was measured in 2-h two-bottle choice (2BC) sessions started 30 min after CNO injection. CNO vs. Vehicle: **, p<0.01, paired t-test.

Accordingly, while it is possible that the small subset of PSTN neurons co-expressing *Crh* and *Tac1* promotes alcohol intake via NK1 signaling, the activity of PSTN *Tac1* neurons as a whole inhibits alcohol drinking. This inhibitory influence of PSTN *Tac1* neurons extends to saccharin drinking and is consistent with the general pattern of feeding/drinking suppression previously reported for this population (40, 43, 44).

### Chemogenetic stimulation of PSTN *Crh* neurons promotes digging, active coping, and exploration, and elevates mechanical pain thresholds

Increases in alcohol consumption can result from various sources of positive and negative reinforcement (5, 45–47). We thus sought to determine the affective and nociceptive state of mice that undergo chemogenetic stimulation of PSTN *Crh* neurons. *Crh*-Cre mice were injected in the PSTN with a Cre-dependent hM3Dq (or mCherry) vector and subjected to a test battery probing their behavior following CNO (or vehicle) administration (Fig. 5A).

In the digging assay (Fig. 5B), there was a significant vector x CNO interaction for the latency to start digging (F_1,22_=9.0, p=0.0065), the number of digging bouts (F_1,22_=13.5, p=0.0013) and the total duration of digging (F_1,22_=11.7, p=0.0025), which reflected a faster onset and robust increase in digging activity in hM3Dq mice (p’s<0.0001), but not in mCherry controls (p’s>0.61), following CNO administration. This effect was confirmed in an independent cohort of hM3Dq mice (Fig. S7A; main effect of CNO for latency: F_1,14_=24.3, p=0.0002; bouts: F_1,14_=33.2, p<0.0001; duration: F_1,14_=17.4, p=0.0010), and no sex differences were observed (sex x CNO interaction for latency: F_1,14_=0.00025, p=0.99; bouts: F_1,14_=1.7, p=0.21; duration: F_1,14_=1.3, p=0.28).

In the tail suspension test (Fig. 5C), there was also a significant vector x CNO interaction for the immobility duration (F_1,22_=9.4, p=0.0057), whereby CNO reduced immobility in hM3Dq mice (p<0.0001), but not in mCherry controls (p=0.57). This effect was confirmed in an independent cohort of hM3Dq mice (Fig. S7B; F_1,14_=90.0, p<0.0001), and no sex differences were observed (F_1,14_=1.5, p=0.24).

In the elevated plus-maze (Fig. 5D), there was a significant vector x CNO interaction for the distance traveled in the closed arms (F_1,20_=5.4, p=0.030) and open proximal arms (F_1,20_=9.5, p=0.0060), reflecting higher locomotion across the maze in CNO-treated hM3Dq mice (closed: p=0.016; open proximal: p=0.0024), but not mCherry controls (p=0.93 and p=0.81, respectively), compared to vehicle-treated counterparts. The interaction was also significant for the number of entries into the proximal (F_1,20_=8.6, p=0.0081) and distal (F_1,20_=4.7, p=0.043) segments of the open arms, as well as for the time spent at the end of the open arms (F_1,19_=8.9, p=0.0077), reflecting increased exploration of the exposed parts of the maze by CNO-treated hM3Dq mice (open proximal entries: p=0.0027; open distal entries: p=0.019; open distal time: p=0.027).

These effects were generally confirmed in an independent cohort of hM3Dq mice (Fig. S7C), in which CNO increased the distance traveled, number of entries, and time spent in the proximal (distance: F_1,20_=26.4, p<0.0001; entries: F_1,20_=9.7, p=0.0056; time: F_1,20_=10.7, p=0.0038) and distal (distance: F_1,20_=26.6, p<0.0001; entries: F_1,20_=30.7, p<0.0001; time: F_1,20_=9.4, p=0.0061) segments of the open arms. In this cohort, CNO tended to increase the distance traveled (F_1,20_=3.9, p=0.061) in the closed arms, an effect driven by the females. In contrast, CNO significantly reduced the time spent in this zone (F_1,20_=13.6, p=0.0014) and this effect was more pronounced in males. Females traveled more distance than males in the closed arms (main effect of sex: F_1,20_=7.9, p=0.011). Conversely, males tended to make more entries (F_1,20_=3.1, p=0.092) and spend more time (F_1,20_=3.2, p=0.088) in the proximal segments of the open arms compared to females. No other sex differences were detected (F’s<2.3, p’s>0.15).

In the tail pressure test (Fig. 5E), there was a significant vector x CNO interaction (F_1,22_=11.6, p=0.0025), whereby CNO elevated the mechanical nociceptive thresholds of hM3Dq mice (p=0.0005), but not mCherry controls (p=0.88).

Overall, we found that the chemogenetic stimulation of PSTN *Crh* neurons is associated with increased digging activity, increased mobility in response to an inescapable stressor (active coping), increased exploration of innately aversive spaces, and reduced pain sensitivity. Taken together, these phenotypes reflect a state of behavioral disinhibition and are not consistent with negative affect.

### CRF synthesis in the PSTN accelerates alcohol drinking escalation and prolongs withdrawal-induced hyperalgesia in the CIE-2BC model

We then asked whether CRF signaling originating from the PSTN might contribute to ethanol intake escalation and other behavioral phenotypes induced by CIE exposure. To do so, we first validated a local knockdown approach by injecting a vector encoding an shRNA targeted against *Crh* (shCrh), or a control shRNA sequence (shControl), in the PSTN of C57BL/6J mice (Fig. 6A). The shCrh vector strongly reduced PSTN *Crh* expression compared to shControl (Fig. 6A; t_5_=8.9, p=0.0003). These vectors were then used in a behavioral cohort.

**Figure 5.**
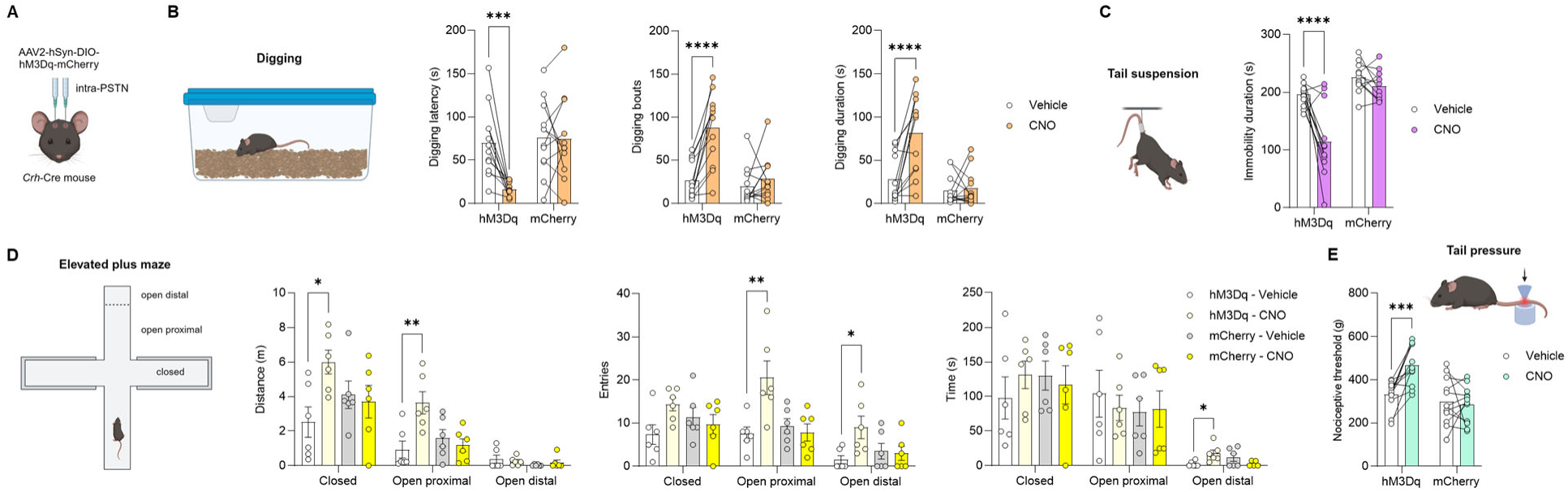
Chemogenetic stimulation of PSTN *Crh* neurons promotes digging, active coping, and exploration, and elevates mechanical pain thresholds. **A.** *Crh*-Cre mice were injected with a Cre-dependent hM3Dq-encoding vector (or mCherry control) in the PSTN. Digging (**B**), tail suspension (**C**), elevated plus-maze (**D**), and tail pressure (**E**) tests were conducted 30 min after CNO administration. CNO vs. Vehicle: *, p<0.05; **, p<0.01; ***, p<0.001; ****, p<0.0001, Šídák’s *posthoc* tests. As expected, CNO had no significant effect in mice expressing mCherry only.

**Figure 6.**
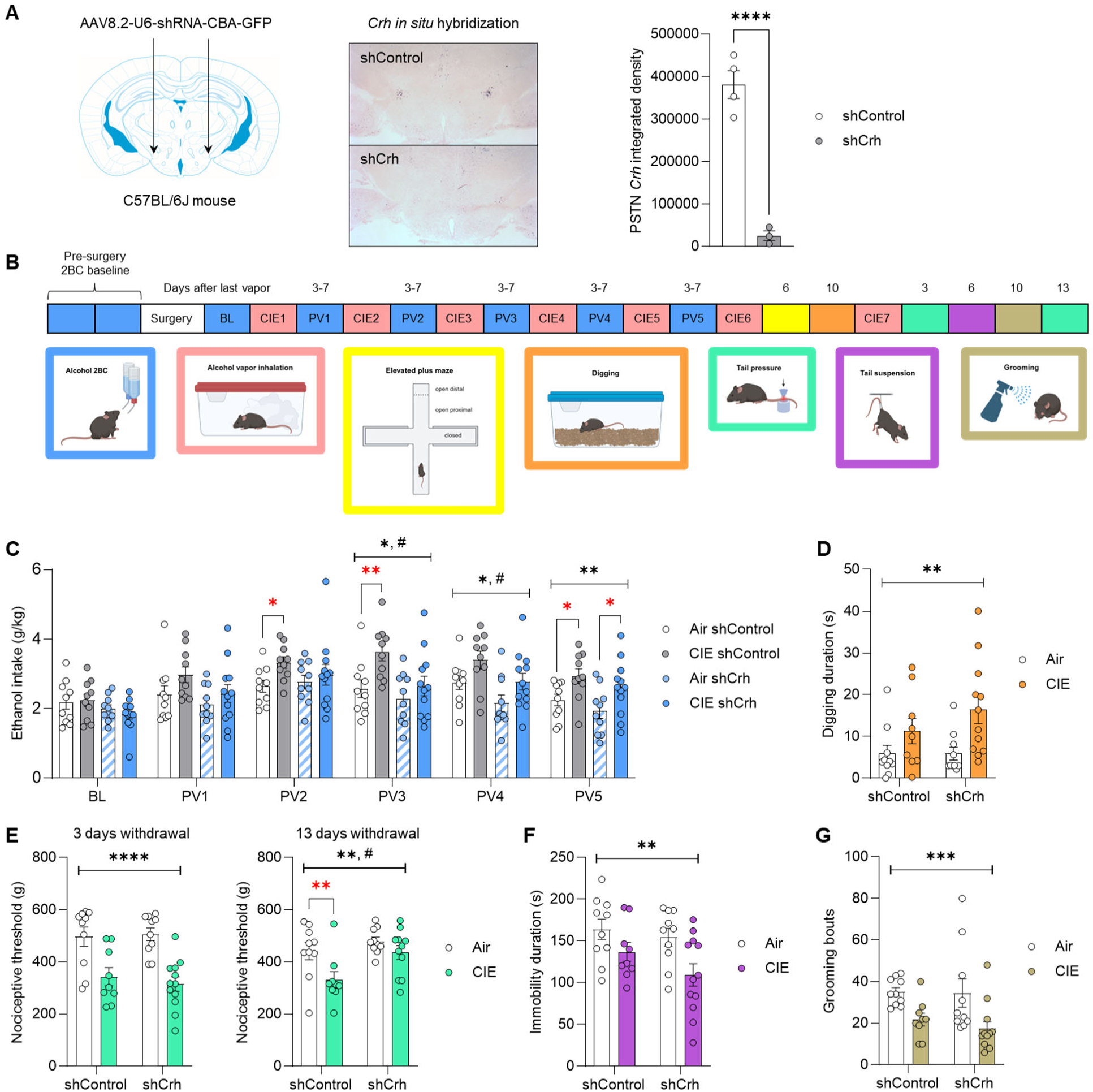
CRF synthesis in the PSTN accelerates alcohol drinking escalation and prolongs withdrawal-induced hyperalgesia in the CIE-2BC model. C57BL/6J mice were injected in the PSTN with a vector encoding an shRNA targeting *Crh* (shCrh) or a control shRNA sequence (shControl). A. *Crh* expression was visualized by chromogenic *in situ* hybridization and signal density in the PSTN was quantified. shCrh vs. shControl: ****, p<0.0001, unpaired t-test. **B.** Another cohort was subjected to behavioral testing. **C.** Ethanol intake was measured at baseline (BL) and after each week of Air/CIE exposure (PVn, post-vapor week n). Digging (**D**), tail pressure (**E**), tail suspension (**F**), and splash (**G**) tests were conducted 3-13 days after last vapor exposure, as shown in panel **B**. Main effect of CIE (black stars): *, p<0.05; **, p<0.01; ***, p<0.001; ****, p<0.0001. Main effect of shRNA: ^#^, p<0.05. Red stars represent unprotected CIE vs. Air comparisons within shControl or shCrh mice: *, p<0.05; **, p<0.01.

C57BL/6J mice were first trained to drink alcohol and split into two groups of equivalent baseline intake assigned to the shCrh or shControl vector. Three weeks after vector injection, alcohol 2BC sessions were resumed and the two groups were further split for assignment to Air or CIE exposure (Fig. 5B). The shCrh vector tended to reduce baseline alcohol intake, prior to CIE exposure (BL; main effect of shRNA: F_1,38_=3.4, p=0.074). A trend for an effect of CIE emerged during the first (PV1; F_1,38_=3.5, p=0.070) and second (PV2; F_1,38_=3.8, p=0.057) post-vapor weeks. On the third (PV3) and fourth (PV4) post-vapor weeks, the main effects of CIE (PV3: F_1,38_=7.3, p=0.010; PV4: F_1,38_=7.0, p=0.012) and shRNA (PV3: F_1,38_=5.8, p=0.021; PV4: F_1,38_=6.6, p=0.014) were both significant, reflecting ethanol intake escalation in CIE mice and reduced intake in shCrh mice. On PV2 and PV3, CIE shControl mice consumed significantly more alcohol than their Air counterparts (PV2, p=0.044; PV3, p=0.0083), while no such difference was observed among shCrh mice (PV2, p=0.52; PV3, p=0.33). By the fifth post-vapor week (PV5), the main effect of CIE was still significant (F_1,38_=10.3, p=0.0027), but the effect of shRNA was no longer significant (F_1,38_=2.4, p=0.13), as both shControl and shCrh mice had escalated their intake in response to CIE (p=0.030 and p=0.030, respectively). Accordingly, *Crh* silencing in the PSTN slows down, but does not prevent, the escalation of alcohol drinking in mice exposed to CIE. The slower rate of escalation was not explained by differential levels of intoxication during vapor inhalation, as there was no significant effect of shRNA on blood alcohol levels measured across CIE weeks (Fig. S7A; F_1,20_=0.81, p=0.38).

The mice were then exposed to additional weeks of CIE to measure affective and nociceptive responses during withdrawal. In the digging assay (Fig. 6D), there was a significant main effect of CIE (F_1,37_=8.9, p=0.0051), reflecting increased digging activity in CIE-withdrawn mice across the two shRNA conditions. There was also a significant main effect of CIE on mechanical nociceptive thresholds measured 3 and 13 days after last vapor exposure (Fig. 6E; day 3: F_1,37_=31.0, p<0.0001; day 13: F_1,36_=7.9, p=0.0078), reflecting CIE-induced hyperalgesia.

However, at withdrawal day 13, the effect of shRNA was also significant (F_1,36_=7.0, p=0.012), and while CIE significantly lowered the threshold of shControl mice (p=0.0082), it did not affect shCrh mice (p=0.26). In the tail suspension test (Fig. 6F), CIE reduced immobility in both groups (F_1,37_=8.7, p=0.0054). Likewise, in the splash test, CIE reduced grooming irrespective of shRNA (F_1,37_=13.0, p=0.0009). There were no significant effects of vapor or shRNA on EPM measures (Fig. S7B-D; F’s<2.9, p>0.10). Altogether, silencing *Crh* expression in the PSTN accelerated the recovery from mechanical hyperalgesia during protracted abstinence, but did not influence the excessive digging, active stress coping, or grooming deficits associated with CIE withdrawal.

## Discussion

Our results demonstrate for the first time that PSTN *Crh* neurons exert bidirectional control over the consumption of fluid reinforcers. Specifically, stimulating PSTN *Crh* neurons is sufficient to robustly increase alcohol and saccharin intake in limited-access free-choice sessions, and this effect does not result from increased thirst. Conversely, inhibiting PSTN *Crh* neurons reverses the escalation of alcohol and saccharin intake produced by CIE exposure. The ability of PSTN *Crh* neurons to increase alcohol drinking is not mediated by CRF_1_ signaling and generally resists pharmacological manipulations that reduce alcohol drinking under control conditions. Furthermore, the behavioral changes elicited by the stimulation of PSTN *Crh* neurons reflect a pattern of disinhibition and pain insensitivity. Finally, we found that silencing *Crh* expression in the PSTN slows down the escalation of alcohol intake in mice exposed to CIE and accelerates their recovery from withdrawal-induced mechanical hyperalgesia.

The PSTN is generally known to suppress consummatory behaviors (see 17 for review). It becomes activated upon sudden food ingestion, exposure to aversive stimuli that reduce feeding (e.g., visceral malaise, novelty, predator odor), and administration of anorectic hormones, thus encoding states of satiety and food rejection. The functional manipulation of PSTN glutamatergic neurons as a whole or subsets of PSTN neurons (e.g., those expressing *Tac1* or *Adcyap1*) demonstrated that their activation serves to suppress food or sucrose intake (40, 43, 44, 48–51). Consistent with prior literature, chemogenetic activation of PSTN *Tac1* neurons virtually ablated the consumption of alcohol and saccharin in our experimental conditions. In this context, our finding that PSTN *Crh* neurons promote the consumption of alcohol and saccharine sets these neurons apart as a unique subset opposing the influence of neighboring populations. In a recent study, we found that activating PSTN *Crh* neurons promotes the consumption of a novel, palatable food (Froot Loops) in hungry mice, and the consumption of a novel, palatable fluid (sucrose) in thirsty mice (44). Here, we show that this effect extends to alcohol and saccharin drinking, does not require fluid deprivation, and withstands the concomitant availability of water (free-choice consumption) and extensive habituation to the reinforcers. The present study represents the first investigation of the PSTN in the context of psychotropic substances and future studies will determine whether its influence on drug self-administration is specific to orally ingested reinforcers or extends to intravenously infused reinforcers.

We used CIE as a well-established experimental modality to increase voluntary alcohol drinking in mice (38). We found that CIE-exposed mice not only consumed more alcohol, but also more saccharin, than their air-exposed counterparts. A previous study had established that CIE does not escalate sucrose consumption in mice that are trained to drink sucrose but are never given access to alcohol drinking (38). Our observation suggests that voluntary consumption of alcohol is required for CIE to induce a concomitant escalation of saccharin consumption. Alternatively, the differential taste and caloric properties of sucrose and saccharin may explain the divergent phenotypes. In any case, it appears that the increased self-administration of alcohol elicited by CIE can generalize to a sweet reinforcer in mice, which may be relevant to the preference for stronger sweet solutions that has been repeatedly observed in humans with an alcohol use disorder (AUD) (52–56) and supports the notion that increased sweet liking might be a consequence of chronic alcohol consumption rather than a predisposing factor (57, 58). The activation of the dorsal anterior insula (which projects to the PSTN in mice (43)) in response to an intensely sweet taste correlates with the enjoyable effect of alcohol (59), suggesting that the association between AUD and sweet liking is mediated by a higher sensitivity to positive reinforcement. Inhibiting PSTN *Crh* neurons reduced both alcohol and saccharin intake and this effect was more pronounced in CIE-exposed mice, supporting the notion that sweet liking exacerbation in AUD shares a common neural substrate with increased alcohol consumption. This finding also suggests that the endogenous activity of PSTN *Crh* neurons is higher during protracted withdrawal from CIE than in air-exposed counterparts. Future studies will test this hypothesis and investigate the molecular changes that may be produced by CIE exposure in this cell type.

Based on the extensive literature implicating excessive CRF_1_ signaling in escalated alcohol consumption in rodents (3, 4, 6), we had hypothesized that the enhanced alcohol drinking driven by PSTN *Crh* neurons may be mediated by CRF release and subsequent activation of CRF_1_ receptors. However, the CRF_1_ antagonist CP376395 was ineffective at lowering alcohol intake following chemogenetic activation of PSTN *Crh* neurons, even though it tended to reduce alcohol consumption in control conditions, in accordance with prior data (12, 26, 60). None of the other ligands we tested selectively reduced the enhancement of alcohol drinking produced by CNO. Most were ineffective at lowering alcohol intake following chemogenetic activation of PSTN *Crh* neurons. Blocking NK1 receptors or mGlu5 receptors significantly reduced alcohol consumption across the vehicle and CNO conditions suggesting that these receptors may participate in the effect of chemogenetic activation. However, alcohol intake was still 2-3x higher after CNO than vehicle injection even in the presence of the highest dose of NK1 or mGlu5 antagonist, indicating that additional pharmacological mechanisms remain to be uncovered. Future research will aim to identify other neurochemicals produced by PSTN *Crh* neurons to guide the selection of pharmacological agents that may succeed in blocking the effect of CNO on alcohol intake. A complementary question relates to localizing the projection(s) that mediate(s) the ability of PSTN *Crh* neurons to increase alcohol drinking.

In addition to increasing alcohol and saccharin consumption, we found that activating PSTN *Crh* neurons reproducibly increased digging activity, mobility in the tail suspension test, and exploration of the exposed arms of an elevated plus-maze. It also reduced pain perception in a mechanical nociception assay. This combination of phenotypes does not support the notion that PSTN *Crh* neuron activity would elicit a state of emotional or physical distress (hyperkatifeia) that would promote alcohol consumption via negative reinforcement. While anxiolytic drugs can reduce digging, spontaneous digging is an innate, species-typical behavior that does not correlate with other indices of anxiety-like behavior and is instead considered an ethologically relevant index of well-being (61, 62). Interestingly, digging is also reliably increased during withdrawal from chronic alcohol exposure in mice, but the translational significance of this phenotype remains to be determined (63, 64). Mobility in the forced swim or tail suspension assays is thought to represent an active coping strategy in response to an acute unescapable stressor, to be contrasted with the passive coping reflected by immobility (65, 66). Alternatively, increased mobility can be interpreted as lack of adaptive learning, whereby switching to immobility favors energy conservation and survival (67). Some studies have observed a correlation between increased mobility in these assays and independent indices of anxiety-like behavior, suggesting that increased mobility could reflect an anxiogenic-like effect (68).

However, in our study, the increased exploration of the open arms and open arm extremities of the elevated plus-maze instead suggests that PSTN *Crh* neuron activity is anxiolytic rather than anxiogenic. Altogether, the behavioral changes elicited by the stimulation of PSTN *Crh* neurons are consistent with a general pattern of behavioral disinhibition, whereby activity is increased in conditions of high arousal (new bedding in the digging assay, inescapable stressor in tail suspension test, novel environment and approach/avoidance conflict in the elevated plus-maze) and pain sensitivity is reduced. This pattern is consistent with the effect of chemogenetic activation of PSTN calretinin neurons (∼90% of PSTN neurons), which increases wakefulness and exploratory behaviors (69). In humans, specific facets of behavioral disinhibition and impulsivity are linked to alcohol use via a common externalizing factor (70–73). Interestingly, both sweet liking exacerbation and high novelty seeking are positively associated with AUD, even more so when combined (74–78). Our results suggest that dysregulation of PSTN *Crh* neurons could represent a mechanism driving both traits and the propensity to consume more alcohol. As such, the PSTN might have functional relevance for the “Reward type” neurobehavioral profile of addiction, which is characterized by higher approach-related behavior and high resting-state connectivity in the Value/Reward, Ventral-Frontoparietal and Salience networks (79).

Knocking down *Crh* expression in the PSTN resulted in delayed alcohol drinking escalation and faster resolution of hyperalgesia in mice withdrawn from CIE. Accordingly, even though CRF_1_ signaling does not mediate the increase in alcohol drinking induced by acute stimulation of PSTN *Crh* neurons, CRF synthesis in the PSTN contributes to the gradual escalation of alcohol drinking induced by CIE. Additionally, even though activating PSTN *Crh* neurons elevates nociceptive thresholds, the production of CRF in the PSTN contributes to maintaining lower nociceptive thresholds during protracted abstinence, further highlighting the functional dissociation between the roles of CRF synthesis in the PSTN and the activity of PSTN *Crh* neurons in alcohol-related behavioral outcomes. This discrepancy may relate to the firing pattern of PSTN *Crh* neurons, such that, under physiological conditions, the role of CRF produced in the PSTN might result from their asynchronous tonic firing and might not be engaged when chemogenetic activation produces synchronous phasic firing. Another possibility is that CRF synthetized in the PSTN might be released at a timepoint preceding the measure of escalated drinking or hyperalgesia (e.g., during early withdrawal from CIE) and that CRF release from PSTN neurons serves to initiate a signaling cascade that results in increased drinking and sustained hyperalgesia at a later timepoint.

On the other hand, silencing *Crh* expression in the PSTN did not affect the other behavioral phenotypes associated with CIE withdrawal, including increased digging and hypermobility in the tail suspension test, even though similar phenotypes are elicited by the chemogenetic stimulation of PSTN *Crh* neurons. Importantly, these behaviors were tested after CIE-induced alcohol intake escalation had developed to a similar extent in shControl and shCrh mice and they were evaluated at a single withdrawal timepoint (as we sought to avoid issues associated with repeated affective testing). It is thus possible that our experimental design lacked the temporal resolution needed to capture a possible involvement of PSTN *Crh* expression in the development or maintenance of CIE-induced phenotypes.

The increased digging, reduced grooming, and mechanical hypersensitivity we observed in CIE-withdrawn C57BL/6J males are consistent with previous reports, although changes are not always reliably detected (8, 21, 64, 80–82). In the tail suspension test, we had also found increased mobility in males withdrawn from CIE for 11 days, but no difference after 19 days (21, 81). Our observation that CIE-withdrawn mice were more active in response to an acute unescapable stressor is consistent with the effect of early or protracted withdrawal from repeated binge drinking (83–85), but opposite to the effect of protracted withdrawal from chronic continuous alcohol drinking (a paradigm in which mice rarely reach intoxication) (86–89). The data presented here thus corroborate the notion that the stress coping strategy of mice withdrawn from alcohol is sensitive to the modality of alcohol exposure and the withdrawal timepoint. An additional factor that modulates the response of mice to alcohol withdrawal is sex. The CIE-2BC experiments reported in the present study employed males only because our primary goal was to assess the effect of PSTN manipulations on escalated ethanol intake and, from what we and other laboratories have found, CIE-induced alcohol intake escalation is more robust in males than females (21, 90, 91). Males and females were included in all other experiments, and we did not detect sex differences in any of the behavioral effects of chemogenetic stimulation of PSTN neurons. Future research will explore whether PSTN neurons might be differentially recruited between sexes, which could explain why CIE-induced alcohol intake escalation is more consistent in males than females.

Altogether, we identified PSTN *Crh* neurons as a neuronal subpopulation that exerts a potent and unique influence on behavior by promoting the voluntary consumption of alcohol and saccharin, while PSTN neurons are otherwise known to suppress consummatory behaviors. PSTN *Crh* neurons are causally implicated in the escalation of alcohol and saccharin intake in the CIE-2BC mouse model of AUD, but the signaling mechanism mediating this effect remains to be uncovered. The pattern of behavioral disinhibition driven by their activation does not support a role of negative reinforcement as a motivational basis for the concomitant increase in alcohol drinking.

## Acknowledgments

We wish to thank Thao Ngyuen, Sophia Zhu, Tanvi Shah, Ellie Petty, Celeste Moreau, and Maya Mehta Maya for their assistance with histological analysis. The schematics in Figures 1, 2, 4, 5, 6, and S7 were created with BioRender.com.

## Data availability

All data supporting the findings of this study are available within the article and its supplementary information file.

## CRediT authorship contribution statement

**Max Kreifeldt**: Investigation. **Agbonlahor Okhuarobo:** Investigation. **Jeffery L Dunning:** Investigation. **Catherine Lopez:** Investigation. **Giovana Macedo:** Investigation. **Harpreet Sidhu**: Investigation. **Candice Contet**: Conceptualization, Funding acquisition, Project administration, Supervision, Investigation, Formal analysis, Visualization, Writing.

