## Supplementary Figures for "Mouse parasubthalamic *Crh* neurons drive alcohol drinking escalation and behavioral disinhibition"

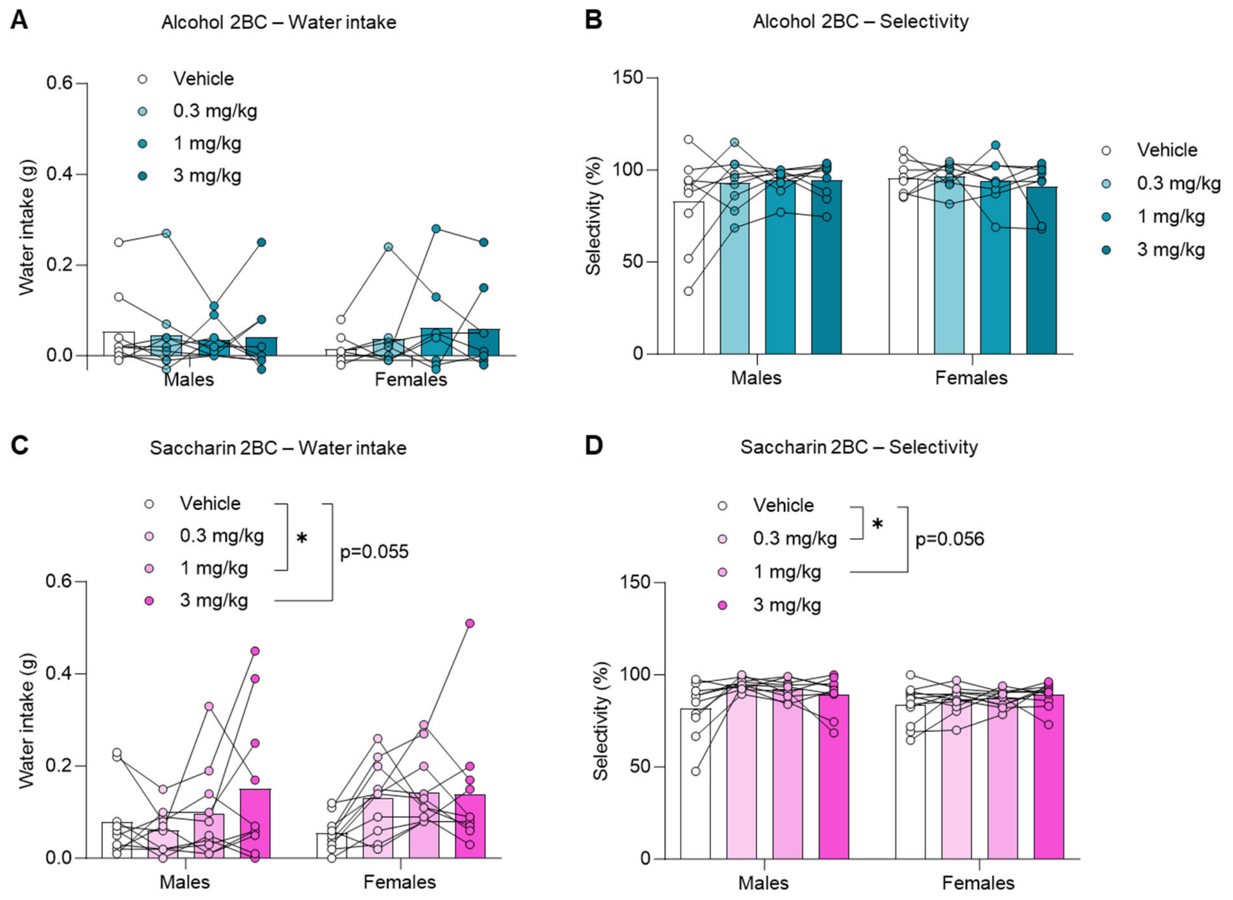


**Supplementary Figure 1.** Water intake (**A, C**) and selectivity (**B, D**) during alcohol (**A-B**) and saccharin (**C-D**) 2BC sessions in Cohort 1. CNO increased water intake and selectivity in the saccharin 2BC assay (*, $p<0.05$, Dunnett's *posthoc* tests).

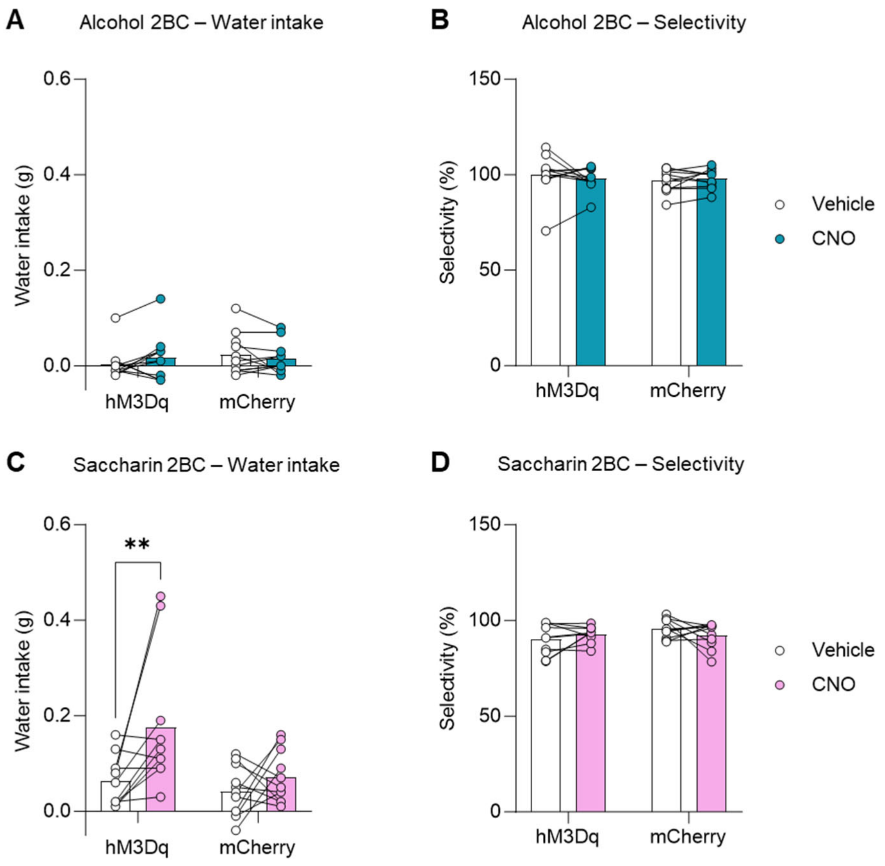


**Supplementary Figure 2.** Water intake (**A, C**) and selectivity (**B, D**) during alcohol (**A-B**) and saccharin (**C-D**) 2BC sessions in Cohort 2. CNO increased water intake in the saccharin 2BC assay (**, $p<0.01$, Šídák's *posthoc* test).

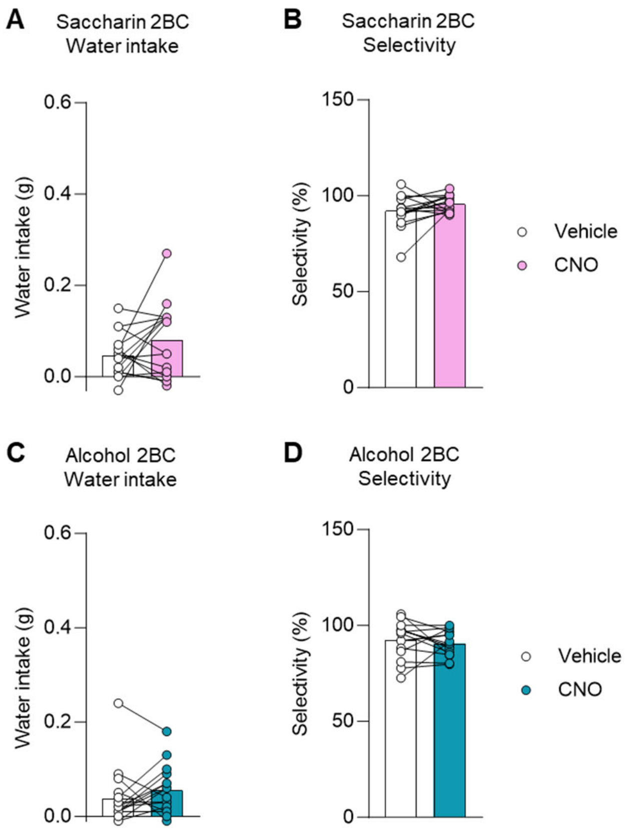


**Supplementary Figure 3.** Water intake (**A, C**) and selectivity (**B, D**) during alcohol (**A-B**) and saccharin (**C-D**) 2BC sessions in Cohort 3. There was no significant effect of CNO on either of these measures (paired t-tests).

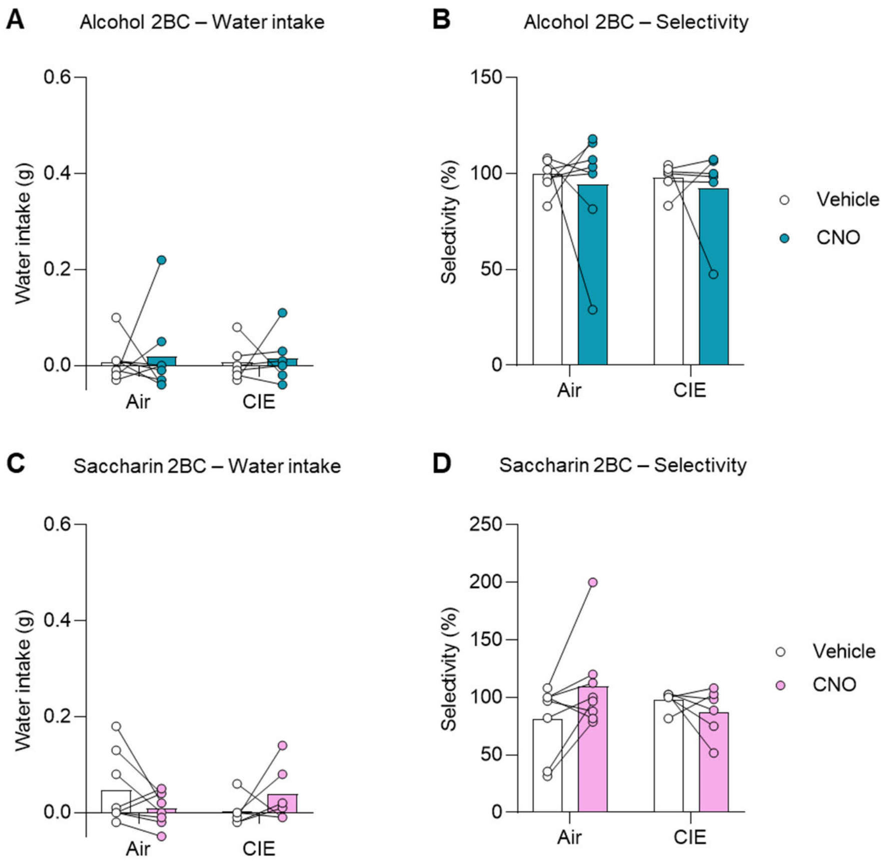


**Supplementary Figure 4.** Water intake (**A, C**) and selectivity (**B, D**) during alcohol (**A-B**) and saccharin (**C-D**) 2BC sessions following chemogenetic inhibition of PSTN *Crh* neurons in CIE-exposed mice. There were no significant main effects of CNO or CIE on either of these measures.

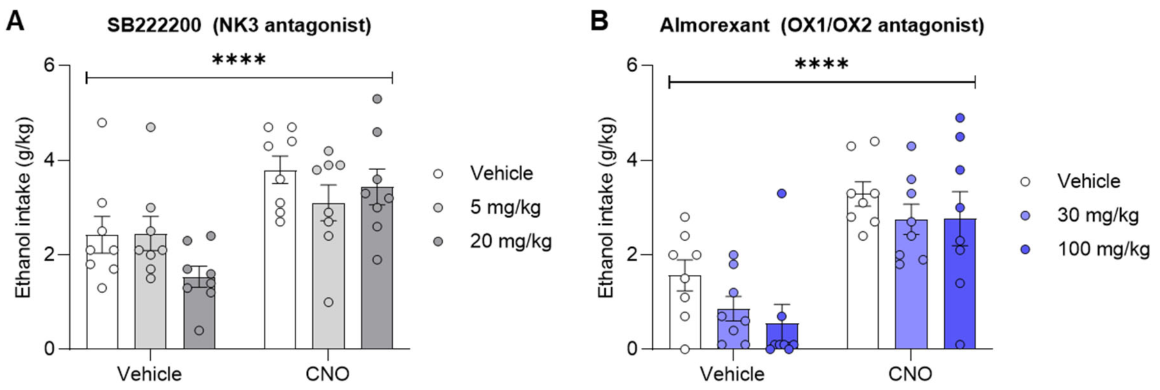


**Supplementary Figure 5.** Ethanol intake following combined administration of SB222200 (**A**) or almorexant (**B**) (between-subjects) and CNO (within-subjects) 30 min prior to 2BC. Main effect of CNO: ****, p<0.0001. The main effect of ligand and CNO x ligand interaction were not significant for either compound.

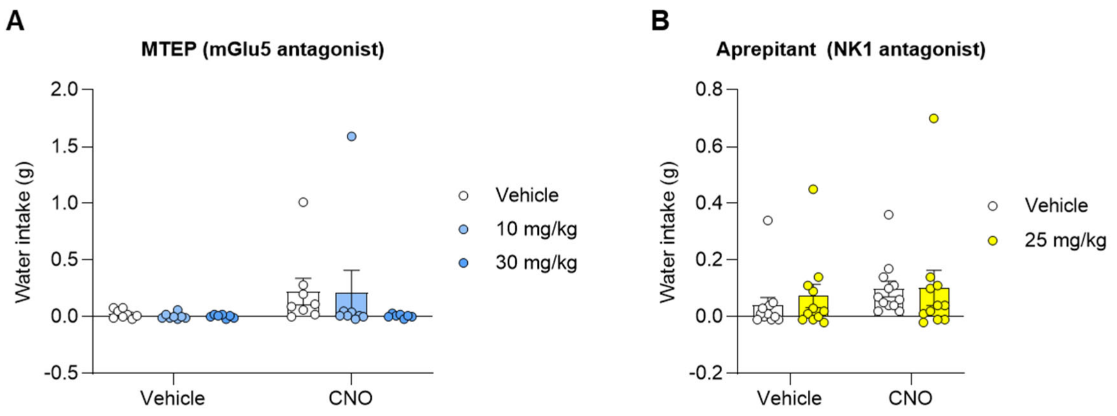


**Supplementary Figure 6.** Water intake following combined administration of MTEP (**A**) or aprepitant (**B**) (between-subjects) and CNO (within-subjects) 30 min prior to alcohol 2BC. The main effects and interaction were not significant for either compound.

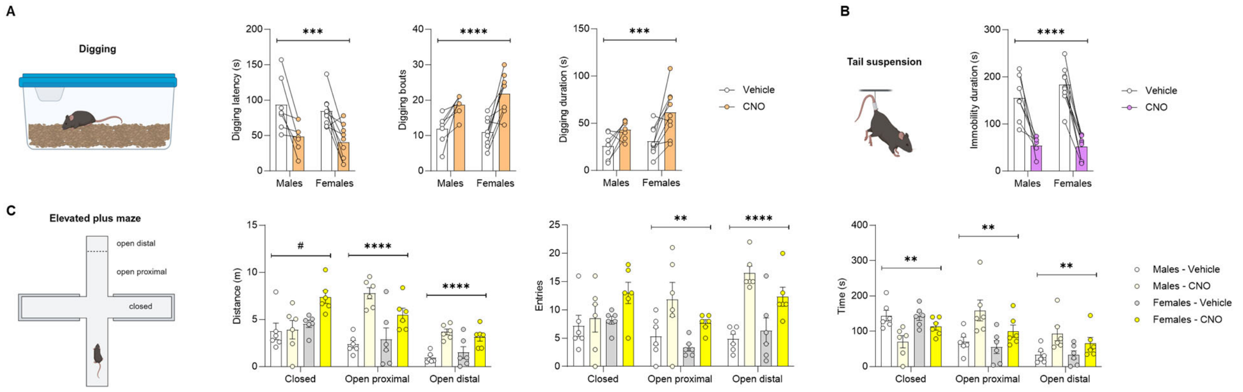


**Supplementary Figure 7.** *Crh*-Cre mice were injected with a Cre-dependent hM3Dq-encoding vector in the PSTN. Digging (**A**), tail suspension (**B**), and elevated plus-maze (**C**) tests were conducted 30 min after CNO administration. Main effect of CNO: *, p<0.05; **, p<0.01; ***, p<0.001; ****, p<0.0001. Main effect of sex: $^{\#}$, p<0.05.

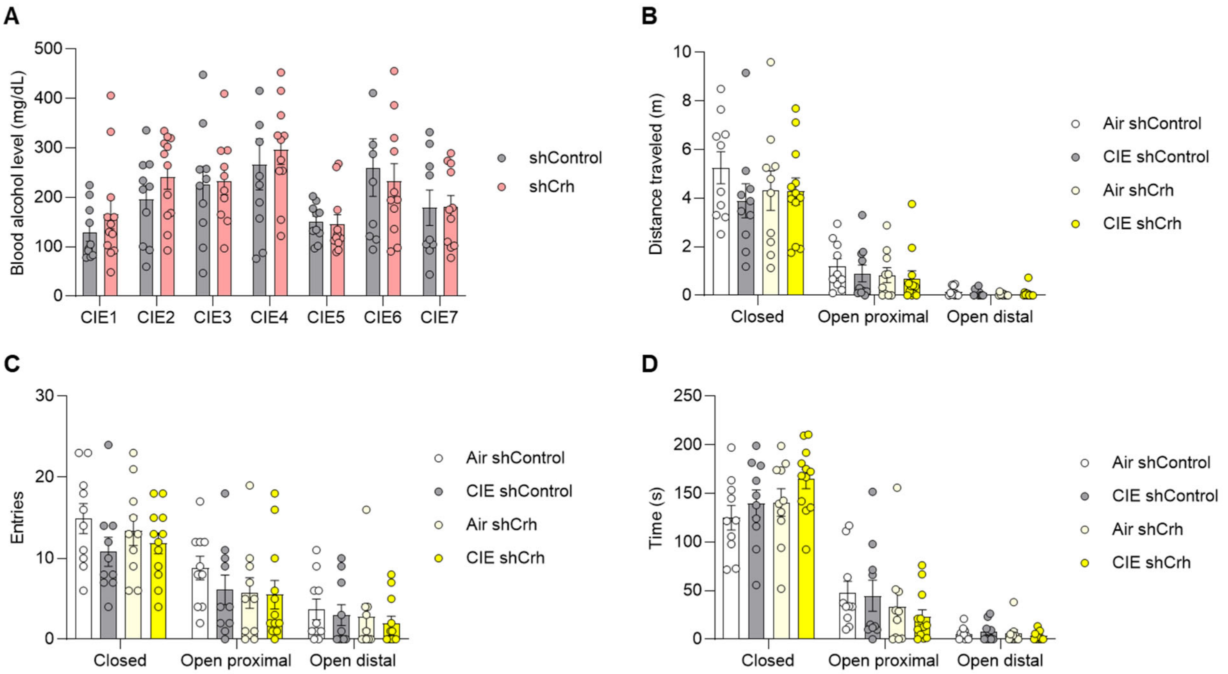


**Supplementary Figure 8.** C57BL/6J mice were injected in the PSTN with a vector encoding an shRNA targeting *Crh* (shCrh) or a control shRNA sequence (shControl) and subjected to the CIE-2BC paradigm. **A.** Blood alcohol levels measured on each week of CIE exposure were similar in the two shRNA groups. **B-D.** Mice were tested in the elevated plus-maze 6 days after CIE week 6. There were no significant effects of vapor or vector on distance (**B**), entries (**C**), or time (**D**) in any of the maze compartments.
